# Strenuous occupational physical activity - potential association with esophageal squamous cell carcinoma risk

**DOI:** 10.1101/469452

**Authors:** Idrees Ayoub Shah, Gulzar Ahmad Bhat, Rumaisa Rafiq, Najma Nissa, Mansha Muzaffar, Malik Tariq Rasool, Abdul Rouf Banday, Mohd Maqbool Lone, Gulam Nabi Lone, Paolo Boffetta, Nazir Ahmad Dar

## Abstract

The impact of leisure time or recreational physical activity (RPA) on cancer risk has been extensively studied. However, the association of the occupational physical activity (OPA), which differs in dose and intensity from RPA, with different cancers including esophageal squamous cell carcinoma (ESCC), has received less attention. Here, we present data from a case-control study which was conducted for assessing the risk factors for ESCC. Histopathologically confirmed 703 ESCC cases and 1664 controls, individually matched to the respective cases for age, sex and district of residence were recruited. Information on type, duration and intensity of physical activity was obtained in face-to-face interviews with participants using a structured questionnaire. Conditional logistic regression models were used to calculate odds ratios (ORs) and 95% confidence intervals (95% CIs). A high level of occupational physical activity was associated with increased ESCC risk (OR = 2.17, 95% CI; 1.41 – 3.32), compared to subjects with moderate occupational physical activity. The association with ESCC risk was stronger in strenuous workers (OR = 3.64, 95% CI; 2.13 – 6.20). The association of strenuous OPA with ESCC risk persisted only in subjects that were involved in strenuous activities for equal to or greater than 5days/week. Our study suggests a possible association of strenuous occupational physical activity with ESCC risk. Although, our results were adjusted for multiple factors including indicators of socioeconomic status, more replicative occupational epidemiological studies are needed to rule out any residual confounding.

## Introduction

Esophageal cancer is the 6^th^ most common cause of cancer related deaths in the world^[1]^. Its two main histological types, esophageal adenocarcinoma (EADC) and esophageal squamous cell carcinoma (ESCC), have different etiology and geographical distribution^[2]^. The incidence of EADC is higher in the developed world, while as ESCC is more common in the developing nations, particularly in Asia, which together contributes about 83% of the total global ESCC burden^[2]^. Unlike EADC^[3]^, the etiology of ESCC is attributed to various indicators of low socioeconomic status (SES)^[4]^ pervasive in the developing nations. Although many such factors including low fruit and vegetable intake^[5, 6]^, poor oral hygiene^[7, 8]^, low SES^[9, 10]^ and frequent and close contact with animals like ruminants^[11, 12]^ have been associated with ESCC risk, the overall etiology of ESCC is not completely understood yet^[13, 14]^. Moreover, unlike low risk regions of ESCC, tobacco and alcohol use are not strongly associated with ESCC risk in these high risk populations^[14]^. Hence, other possible risk factors in these high incidence regions of ESCC need to be explored.

Compared to the developed world, developing economies lack potential industrialization and have usually low technological advantages. As a consequence, most of the occupations like farming, shoveling, heavy weight lifting and construction works etc., involve human exertions and demand excessive physical labor. Such strenuous and exhaustive occupational physical activities (OPA) involved in such jobs are done for many hours a day, for most of the days per week, most often carried out for many years; and thus cannot be compared with recreational physical activity (RPA). Hence, OPA in developing world offers a different setting to study its association with cancer risk. Moreover, the perception is now building that there is a threshold line up to which physical activity can be beneficial. Cuople of recent studies with non connonical findings: One, Million Woman Study^16^ reported that strenuous RPA can increase breast cancer risk and sencond study reproted that more occupational physical activity time increases ESCC risk^[15]^.

ESCC is the most common cancer in Kashmir, India^[16]^, which is under economic transition. Majority of the population live in rural areas, which contributes to the maximum number of ESCC cases^[17]^. The rural inhabitants are commonly engaged in farming activities and other unskilled jobs that by nature involve strenuous physical activity, while as the jobs that involve fewer efforts like government services and business are less common^[9]^. We conducted a case-control study and reported association of number of risk factors in study population with ESCC risk^[8, 11, 18–21]^. Earlier we reported, the people involved in active or very active OPA have higher risk of ESCC, as compared to sedentary life behavior^[9]^. Here we further analyze the data on OPA to investigate the association between level and frequency of OPA with ESCC risk in Kashmir.

## Materials and methods

### Subject selection

We conducted a case control study to identify the various risk factors of the ESCC in Kashmir whose details of the study design and related information has been reported previously^[8, 9, 11, 21]^. Briefly, all histopathologically confirmed ESCC cases (n=703), who had no previous cancer, were recruited from the Regional Cancer Centre and Department of Radiation Oncology of Sher-i-Kashmir Institute of Medical Sciences (SKIMS) from 2008 to 2012. For each case at least one control (n = 1664) individually matched to the sex, age (± 5 years), and district of residence was recruited from in-patient wards of SKIMS, Government Medical College Hospital, Srinagar, and ten district hospitals. Patients were enrolled as controls only when the disease for which they had been admitted did not have a strong association with tobacco or alcohol consumption. The participation rate for cases and controls was 96% (732 invited, 29 refusals) and 98% (1697 invited, 34 refusals) respectively. The majority of those who refused were too ill to participate in the study. We were able to recruit only one control for 45 cases, 2 controls for 377 cases, 3 controls for 269 cases and more than 3 controls for 13 cases. Informed consent was obtained from all subjects. This study was reviewed and approved by the Institutional Ethics Committee of SKIMS.

### Data collection

Structured questionnaires were administered in face-to-face interviews at hospitals by trained interviewers in primary language. A limited number of staff conducted the interviews and no proxies were used. Detailed information on demographic characteristics, habits, including lifelong history for use of alcohol, several tobacco products, and intake of fresh fruits and vegetables was collected. Ever use of alcohol and tobacco products was defined as the use of the respective product at least weekly for a period of 6 months or more. To assess the SES, information on potential parameters of SES was obtained including education level (highest level attained), occupation, professional work intensity, income, house type, cooking fuel, place of residence, and ownership of several household appliances, including personal automobile, motorbike, B/W TV, color TV, refrigerator, washing machine, vacuum cleaner, computer, and bath in the residence. The detailed information about consumption of the various fruits and vegetables was collected using a semi-quantitative food frequency questionnaire. We asked about the usual frequency of use in a day, week, or month and the amount of use in each instance. Using this information, we calculated the intake of each fruit and vegetable in g/day. In order to cover the intake of seasonal fruits/vegetables, we also collected data on the number of months in which any of these fruits were consumed: the daily intake in these cases was multiplied by the number of months of consumption and divided by 12. The daily intake of all fruits and vegetables were summed up to estimate the total fruit/vegetable intake per day.

As the study was an epidemiological study (rather an occupational epidemiological study), we collected information on the type of occupations and nature of the physical activity besides on many potential risk factors prevailing in Kashmiri population. Participants who were involved in the occupations (like shoveling farming, brick or stone setters, landscape works, loggers, construction workers etc.) where strenuous physical activity was possible, were further asked that if the strenuous physical activity was a part of their daily activities. As an indicators of strenuous activity, the subjects where asked if they sweated a lot and heart rate increased during their working time^[22]^. Information on frequency of work done in a week (number of days they work in a week) by the subject was recorded as: A) 1- 2 days in week; B) 3-4 days in week; C) ≥ 5days/week. The information on RPA was also obtained.

### Statistical Analysis

Numbers and percentages were calculated and presented for categorical variables, as well as means and standard deviations (SD) or median and inter-quartile range for continuous variables. Fruit and vegetable intake data (g/day) were transformed to logarithmic values following addition of 0.1 to original values. To assess the SES, we built a composite score for wealth, based on appliances ownership and other variables by using multiple correspondence analysis (MCA)^[10]^. Information on MCA method is provided elsewhere^10^. Conditional logistic regression was used to calculate unadjusted and adjusted odds ratios (ORs) and corresponding 95% confidence intervals (CIs). By design, case and control subjects were matched by age, sex, and district of residence. Adjusted ORs (95% CIs) were obtained from two models. In the first model, ORs (95% CIs) were adjusted for demographic factors, including age, ethnicity, place of residence (rural/urban), religion, and education level. Age was included in the multivariate models, because the matching for age was not perfect (±5 years). We adjusted the results for religion because an earlier study from this region had suggested dissimilar incidence of ESCC among people with different religions^[23]^. In the second group of models, in addition to above demographic factors, some biologic factors, including daily fresh fruit and vegetable intake (logarithmic scale), cumulative use of cigarette, hookah, and *nass*, and ever-use of *bidi, gutka*, and alcohol, were included one by one and then collectively. The occupational physical activities were categorized into two groups; one, which include the participants with least physical activity in their jobs (occupations involving minimum exertions as jobs of clerk, engineer, teacher, business man etc.) and two, very active workers (occupations like shoveling farming, brick or stone setters, landscape works, loggers, construction workers etc.). Further among the active works, those who reported a lot of sweating and substantial increase in heart rate during working hours were grouped as strenuous occupational workers. The number of subjects with 1 – 2 days or 3 – 4 days of work/week was modest, so were grouped together and then participants were categorized into two categories; first, with ≤ 4 days a week and second with ≥ 5days of work/week. All statistical analysis was done using Stata software, version 12 (Stata Corp., College Station, TX, USA). Two sided *P* values < 0.05 were considered statistically significant.

## Results

A total of 703 ESCC cases and 1664 matched controls were recruited in this study. Distribution of demographic factors, socioeconomic indicators and tobacco and alcohol use by case status are shown in Table 1. The mean age of cases and controls was 61.6 and 59.8 years, respectively and ~55% were males. Majority of ESCC cases resided in rural areas (95%). More cases were involved in occupations, which by nature involve strenuous physical activities than the respective controls, while as the jobs like government services and business were more in controls (17%) than the cases (6%). Sizable proportion of ESCC cases was active workers (88%) than controls (64%). The percentage of cases (38%) with strenuous OPA was higher than respective controls (20%). Larger number of cases was smokers and had low formal education than their respective controls. The fresh fruit and vegetable consumption and SES was higher in controls than respective cases. None of the participants reported of RPA.

**Table 1.** Characteristics of 703 ESCC cases and 1664 controls from Kashmir Valley, India, 2008–2012^a^

| **Characteristics** | **Cases (%)** | **Controls (%)** | **P value** |
| --- | --- | --- | --- |
| **Age**, mean (SD), years | 61.6 (11.1) | 59.8 (11.1) |  |
| **Sex** |  |  | 0.78 |
| Men | 393 (55.9) | 920 (55.3) |  |
| Women | 310 (44.1) | 744 (44.7) |  |
| **Place of residence** |  |  | <0.001 |
| Urban | 29 (4.1) | 146 (8.8) |  |
| Rural | 674 (95.9) | 1518 (91.2) |  |
| **Ethnicity** |  |  | 0.58 |
| Kashmiri | 682 (97.0) | 1619 (97.3) |  |
| Gojri | 11 (1.6) | 16 (1.0) |  |
| Pahari | 9 (1.3) | 27 (1.6) |  |
| Other | 1 (0.1) | 2 (0.1) |  |
| **Religion** |  |  | 0.03 |
| Muslim | 695(98.9) | 1648(99.0) |  |
| Hindu | 5 (0.7) | 2 (0.1) |  |
| Sikh | 3 (0.4) | 14 (0.8) |  |
| **Fresh fruit and vegetable intake**, median g/day (IQR) | 7.9 (3.8 – 12.6) | 25.2 (12.0-60.9) | <0.001 |
| **Hookah smoking** |  |  | <0.001 |
| Never | 282 (40.2) | 964 (58.0) |  |
| 1 – 139 hookah-years | 97 (13.8) | 228 (13.7) |  |
| 140 – 240 | 110 (15.7) | 245 (14.8) |  |
| > 240 | 213 (30.3) | 224 (13.5) |  |
| **Cigarette smoking** |  |  | 0.01 |
| Never | 632 (90.0) | 1437 (86.4) |  |
| 1 – 6.2 pack-years | 23 (3.3) | 77 (4.6) |  |
| 6.3 – 13.1 | 21 (3.0) | 73(4.4) |  |
| ≥ 13.2 | 26 (3.7) | 76 (4.6) |  |
| **Bidi ever smoking** | 15 (2.1) | 3 (0.2) | <0.001 |
| **Nass chewing** |  |  | <0.001 |
| Never | 501 (71.6) | 1471 (88.5) |  |
| 1 – 119 *nass*-years | 46 (5.6) | 52 (3.1) |  |
| 120 – 199 | 36 (5.1) | 71 (4.3) |  |
| ≥ 200 | 117 (16.7) | 69 (4.1) |  |
| **Gutka ever chewing** | 10 (1.4) | 13 (0.8) | 0.01 |
| **Alcohol ever use** | 8 (1.1) | 0 (0.0) | <0.001 |
| **Animal contact** |  |  | <0.001 |
| No or occasional contact | 164 (23.3) | 774 (46.5) |  |
| Daily contact | 175 (24.9) | 616 (37.0) |  |
| Daily and close contact | 364 (51.8) | 274 (16.5) |  |
| **Second hand smoking** |  |  |  |
| No second hand smoke exposure | 642 (91.4) | 1504 (93.9) |  |
| Exposure to second hand smoke | 61 (8.6) | 97 (6.1) |  |
| **Family history of cancer** |  |  | <0.001 |
| Negative history | 462 (65.7) | 1544 (92.8) |  |
| Positive history | 241 (34.3) | 120 (7.2) |  |
| **Oral Hygiene** |  |  | <0.001 |
| Don't brush | 161 (22.9) | 101 (6.2) |  |
| Ones/week | 366 (52.1) | 785 (47.9) |  |
| Twice or thrice/week | 94 (13.4) | 345 (21.1) |  |
| Daily | 81 (11.6) | 405 (24.8) |  |
| **Tea type** |  |  | 0.041 |
| Other | 09 (1.3) | 51 (3.1) |  |
| Salt tea | 693 (98.7) | 1612 (96.9) |  |
| **Monthly income(INR)** |  |  | <0.001 |
| 1000-5000 | 514 (77.0) | 988 (59.5) |  |
| 6000-10,000 | 102 (14.5) | 384 (23.1) |  |
| > 10,000 | 60 (8.5) | 290 (17.4) |  |
| **Wealth Score** |  |  | <0.001 |
| Quintile 1-lowest | 397 (56.5) | 337 (20.2) |  |
| Quintile 2 | 112 (15.9) | 328 (19.7) |  |
| Quintile 3 | 66 (9.4) | 334 (20.1) |  |
| Quintile 4 | 70 (10.0) | 333 (20.0) |  |

<sup>a</sup> Although cases and controls were individually matched, the percentages of cases and controls are not necessarily equal in each sex category, because some cases have one matched control and others have more controls. Numbers may not add up to the total numbers due to missing data in some variables. P values calculated using chi-squared tests for categorical variables (chi-square for trend in variables with more than two categories) and Wilcoxon Rank Sum tests for continuous variables.

An increased risk of ESCC was observed in very active workers (OR= 2.17, 95% CI; 1.41 – 3.32) as compared to workers with moderate activity. On further stratification of active group into strenuous and non-strenuous workers, the association was stronger in strenuous workers (OR=3.64, 95% CI; 2.13 – 6.20), but did not persist in non-strenuous workers. The strenuous activity effect on ESCC risk persisted only in subjects who did the strenuous ≥ 5days/week (OR = 3.52, 95% CI 1.97 – 6.31), (Table 2). We assessed the differences in the ESCC risk associated with the intensity and frequency of physical activity as determined by gender; the risk persisted only in males (3.39, 95% CI; 1.96 – 5.88), but was lost in females, may be because of the low number in the model (Supplementary table S1). We further classified the subjects into two groups: one group included farmers, while as other group included non-farming professionals. The ESCC risk was retained in both groups, albeit with a wider CI’s in the non–farmer group, possibly due to low number in the latter subgroup. On further subclassification of the subjects on the basis of cumulative wealth score and monthly income, we observed high ESCC risk in the subjects with strenuous physical activity and low wealth score and this risk was almost significant in participants with high wealth score as well (Supplementary table S2). The outcome of the study did not change when adjustment was done for socioeconomic indicators, education, income and cumulative wealth score one by one (Supplementary table S3).

**Table 2.** Association between physical activity and ESCC risk, Kashmir Valley, India.

| Physical activity | Cases (%) | Controls (%) | Unadjusted OR *<br>(95% CI) | Adjusted OR <sup>a</sup><br>(95% CI) | Adjusted OR <sup>b</sup><br>(95% CI) |
| --- | --- | --- | --- | --- | --- |
| **Occupational physical activity** |  |  |  |  |  |
| Sedentary/ moderate activity | 536 (76.2) | 1506 (90.6) | Referent | Referent | Referent |
| High activity | 167(23.8) | 157 (9.4) | 3.32 (2.52 – 4.36) | 2.53 (1.87 – 3.43) | 2.17 (1.41 – 3.32) |
| **Intensity of work** |  |  |  |  |  |
| Non strenuous | 41 (5.8) | 83 (5.0) | 1.48 (0.98 – 2.23) | 1.18 (0.75 – 1.84) | 1.11 (0.57 – 1.72) |
| Strenuous | 126 (17.9) | 74 (4.6) | 5.63 (3.97 – 7.99) | 4.19 (2.87 – 6.13) | 3.64 ( 2.13 – 6.20) |
| **Frequency of Strenuous work** |  |  |  |  |  |
| ≥ 4 days/week | 6 (0.9) | 7 (0.4) | 2.74 (0.76 – 9.92) | 2.16 (0.43 – 10.98) | 2.29 (0.27 – 19.58) |
| ≤ 5 days/ week | 120 (18.1) | 67 (4.2) | 5.80 (3.98 – 8.44) | 4.23 (2.82 – 6.34) | 3.52 (1.97 – 6.31) |
| *P* for trend |  |  | < 0.001 | < 0.001 | < 0.001 |

Numbers may not add up to the total numbers due to missing data in some variables.
\* By design, controls were individually matched to cases for age, sex, and district of residence.
<sup>a</sup> Adjusted for age, ethnicity, place of residence, religion, and education.
<sup>b</sup> Adjusted for age, ethnicity, place of residence, religion, education, daily fresh fruit and vegetable intake (logarithmic scale), wealth score , monthly income cumulative use of cigarette, hookah, and nass, and ever-use of bidi, *gutka*, and alcohol, exposure to passive smoke, family history of any cancer, oral hygiene, consumption of salt tea, contact with animals.

## Discussion

Our study identified an association between occupational physical activity and ESCC risk. The risk was stronger in strenuous workers, which was retained in men in a dose dependent manner. The risk of ESCC was independent of SES and did not persist in non-strenuous workers; suggesting that this association in very active group is the proxy of strenuous OPA. Previous epidemiological studies on the association RPA with different cancers have reported, in a fairly consistent manner, an inverse association with cancer risk [24–28]. In relation with esophageal cancer, most of these studies are either from the developed world or on EADC. The risk reduction observed in case of EADC amongst the most physically active people compared to the least active people has been attributed to the reduction in obesity-associated chronic inflammation^[29]^, which plays a critical role in the etiology of EADC^[30]^. Conversely, reports on the association of OPA with different cancers have shown mixed results. It has been positively associated with gastric cancer^[31]^ and lung cancer^[32]^, but inversely associated with bladder cancer^[33]^ and EADC [29, 34–36]. Further, in relation with ESCC, three studies are available on OPA; one has reported no association^[35]^, while as the other reported an inverse association^[37]^ in females only. The later study^[37]^, unlike ours, did not stratify the subjects into strenuous and non-strenuous workers and was conducted on relatively smaller sample size. While as the 3^rd^ one has [15] has reported increased OPA time increases risk of ESCC. As of now, no study has tested the association of ESCC risk with the different levels of strenuousness of OPA. More over the findings from Million Woman Study [16], The Copenhagen City Heart Study’^[37, 38]^ and the OPA time^[15]^ have strengthened the notion which propounds upon, that a line needs to be drawn up to which even the strenuous RPA can be beneficial; reporting that even strenuous RPA is associated with an increased vascular risk.

In the current study, the risk of ESCC increased with the level of OPA. It will be unreasonable to extend the protective effect of RPA at least through inflammation reduction^[25]^ to strenuous workers, who get more exhaustion for longer times and are unlikely overweight^[38]^. Interestingly, none of the participants recruited in the study reported any RPA and thus makes this study unique in its ability to detect an effect of OPA. In the current study the association is unlikely a manifestation of any known risk factor of ESCC, as we have adjusted extensively with many such confounders.

To explain the biology of the association of ESCC risk with strenuous occupational activities, one of the plausible mechanisms can be linked to high energy demand that warrants excessive aerobic metabolism. Muscle oxygen uptake and utilization increases up to 200 times during the strenuous exercise^[39–42]^. As an outcome, flux of electron through the rapidly respiring mitochondria in the active muscle appreciably increase the possibility of electron leakage and consequent reactive oxygen species (ROS) production, which is over and above the capacity of the antioxidant scavenging system of the cell^[43]^. Hence during strenuous work the oxidative stress is caused as a result of enhanced production of ROS and free radicals^[39, 43, 44]^.

ROS react with cellular components, carbohydrates, proteins, lipids and DNA and results into cellular or tissue injury, the finger prints of oxidative stress driven carcinogenesis^[45–48]^. For example, DNA oxidation by these reactive species generates 8-hydroxy-2′-deoxyguanosine, a product able to generate mutations in DNA, eventually leading to the development of cancers^[49]^. ROS can also cause DNA damage and mutations via lipid peroxidation. The byproduct of lipid peroxidation (Malondialdehyde, and malondialdehyde - DNA adducts (M1-dG), have been reported in higher levels in tumor than normal human esophageal tissue^[50]^, suggesting that M1-dG adducts may be involved in the initiation or progression of cancer. Further, our argument regarding the enhanced risk of ROS linked to strenuous OPA is supported by various recent reports which have associated oxidative stress with ESCC^[51]^, oral squamous cell carcinoma^[52, 53]^, gastric^[54]^, breast^[55]^, lung^[56]^ and pancreatic cancers^[57]^. The dose dependent association of the severity of the OPA with ESCC is in line with our argument of enhanced ROS production during higher working loads on muscles.

Some ROS initiated reactions that escape enzymatic degradation are normally terminated by chain-breaking antioxidants, including water-soluble ascorbate, vitamin E, ubiquinone, and *β*-carotene^[58]^. Fresh fruits and vegetables are the good sources of such antioxidants^[59]^. In the current study, the fresh fruit and vegetable consumption was higher in controls than respective cases. So high oxidative stress on one hand and low antioxidant supply to the body on the other is more likely to compound the problem.

Males are more associated with the high energy demanding jobs than females, who are engaged with non-strenuous household activities. Further, we also had a very low representation of female strenuous workers in the current study, and the risk associated with strenuous OPA was retained only in males; and thus could not be compared with females. However, if the association is causal, this observation can be a contributing link for the male dominance in ESCC cases in general; but further research in this direction is warranted to understand the exact biology behind.

Unlike adenocarcinoma of the esophagus, ESSC is considered to be the disease of poor. People who have low SES are more likely to be involved with the jobs demanding strenuous physical activity. Low SES has shown some association with ESCC in this population^10^. To exclude the possibility of residual confounding, we adjusted the results for several SES indicators alone or in combination, however association of ESCC with strenuous OPA persisted in each group and the effect of indicators of SES did not affect outcome of the study. Moreover, on stratification of the subjects on the basis of wealth score we observed that ESCC risk persisted in the subjects irrespective of the wealth score. These observations support our argument that association of strenuous occupational physical activity with ESCC risk is independent of wealth score.

The maximum representation of ESCC cases in the current study was from rural areas, where mostly people are occupied with farming and related occupations. On classifying the subjects into farmers and other profession (other group included all non-farming occupations), the ESCC risk persisted in the subjects that were used to strenuous works in both groups.

Histological verification of ESCC and adjustments of the results for multiple potential confounding factors and SES indicators are the major strengths of this study. Information on strenuous physical activity provided by a subject is unlikely to be biased, as job or occupation are less forgettable. Further strenuous physical activity is not an established risk factors for ESCC, so it is unlikely that participants would preferentially misreport activity level at work. Similar to other case-control studies with retrospective exposure assessments, recall and interviewer bias may be also be a concern in this study. However a limited staff interviewed the participants, and nature of job/occupation of a subject is less forgettable. In addition the results in the adjusted model were not adjusted with body mass index (BMI) as the data for BMI was not available. Many important variables with regards to association with OPA were either not collected or not adjusted for. These include height, weight, body mass index, factors associated with health, disease, and physical fitness, exposures to occupational hazards, etc. Further this case-control study was not designed as an occupational epidemiologic study, comprehensive information was not obtained related to OPA which might have impacted outcome. The cases and controls were not matched on occupation and can result into a selection bias and seasonal variation in job and other activities were not considered. Other information on the indicators that influence sweating and heart rate are not available. The details of total physical activity (non-occupation related such as from household chores, etc.) are missing. The study needs to be reproduced with working on the above mentioned limitations.

In summary, the study showed that strenuous occupational physical activity is associated with the increased risk of ESCC in a dose dependent manner in males. However, this association needs validation by further occupational epidemiological studies.

## Acknowledgement

This study was supported by Extramural grant of Indian Council of Medical Research (ICMR), New Delhi, India (File No. 5/13/37/2007/-NCD-III), and Department of Science and Technology, Govt. of India (File No.: SR/SO/HS-07/2009). Idrees Ayoub Shah was awarded Senior Research Fellowship by ICMR vide file No. 3/2/2137/2012/NCD-III. The authors also thank all the participants for volunteering in the study.

